# *In vitro* investigation and evaluation of the antidiabetic potential of the ethanolic extract of *Asparagus racemosus* using starch digestion, glucose diffusion, glucose uptake, and DPPH assays

**DOI:** 10.64898/2026.03.22.713478

**Authors:** Md. Sifat Rahman, J.M.A. Hannan, Rafia Tasnim, Md. Mashrur Meraj Bhuiyan, Chinmoy Basu, Sakibul Haque Sammo, Bijoy Chandra Sarkar, SM Tajul Islam, Sanjid khan

## Abstract

*Asparagus racemosus* commonly known as Shatamull, is a medicinal plant with pharmacological applications documented in both Indian and British Pharmacopoeias and various traditional medicinal practices. Previous studies have reported that *A. racemosus* reduces hyperglycemia by enhancing insulin secretion. The aim of the current study was to assess the antihyperglycemic actions and explore the underlying mechanisms of action of *A. racemosus* utilizing *in vitro* carbohydrate digestion, glucose diffusion, glucose uptake, 2,2-Diphenyl-1-picrylhydrazyl (DPPH) and preliminary phytochemical screening. The inhibition of carbohydrate digestion was assessed using α-amylase and α-glucosidase enzyme assays. The effect on glucose diffusion was evaluated using cellulose ester dialysis tube. Subsequently, glucose uptake was measured in a yeast cell model at different glucose concentrations, and the antioxidant potential was evaluated by measuring DPPH radical scavenging activity. *A. racemosus* significantly reduced (p<0.05– 0.01) glucose release during in vitro starch digestion by 21.11%, whereas glucose absorption decreased by 22.75% (p<0.05–0.001). The most significant enhancement (p<0.05–0.001) in glucose uptake by 65.03%, was observed at 5 mM glucose concentration. Furthermore, it showed significant antioxidant activity by scavenging DPPH (p<0.05, 0.001) radicals by 42.46%. Preliminary phytoconstituent screening indicated the existence of flavonoids, tannins, steroids, glycosides and saponins. In conclusion, *A. racemosus* shows an inhibitory effect on carbohydrate digestion and absorption, enhances glucose uptake and demonstrates significant DPPH radical scavenging activity, potentially due to the presence of naturally occurring phytochemicals. Thus, *A. racemosus* may contribute as a promising antidiabetic drug for the treatment of diabetes mellitus. More investigations are needed to determine the active compounds in *A. racemosus* that contribute to its antidiabetic effects.

## Introduction

Diabetes mellitus is a prevalent metabolic illness marked by sustained hyperglycemia that poses a significant worldwide health issue, involving a myriad of issues affecting multiple organ systems and leading to elevated death and disability rates [1]. Globally, approximately 589 million adults are currently living with diabetes, and around 252 million remain undiagnosed, significantly increasing their risk of serious complications and early mortality. Alarmingly, projections suggest that this number may rise to 853 million by the year 2050 [2]. The fundamental objective of diabetes management is to successfully regulate increased blood glucose levels. The present treatments mainly focus on mitigating postmeal hyperglycemia by limiting the degradation of nutritional starch by the hydrolysis of carbohydrate enzymes such as α-amylase and α-glucosidase, and also by increasing or inducing insulin activity in desired areas through the consumption of antidiabetic drugs [3, 4]. Currently, multiple classes of medicines exist that mimic the functions of insulin by regulating blood glucose levels, including biguanides, sulfonylureas, alpha-glucosidase inhibitors, PPAR-gamma agonists, and SGLT2 inhibitors, which represent the main classes of anti-hyperglycemic medications [5]. Despite that, these synthetically produced antihyperglycemic drugs frequently have serious negative interactions, resulting in an increased mortality risk in the population with diabetes mellitus [6]. Thus, there is an increasing demand for the progress of innovative anti-diabetic medications that improve therapeutic efficacy and reduce toxicity [7].

Recent decades have witnessed an increasing trend in the prospective benefits of plant-based substances for diabetes management because of their efficacy, safety, and minimal adverse reactions [8]. The traditional application of herbal remedies for treating multiple diseases has been practiced for centuries, since plants contain an abundance of pharmacologically potent bioactive compounds with minimal adverse responses. Over 800 types of plants have been found to have antidiabetic properties, and several more are currently under research [9]. These herbal plants contain a wide range of phytoconstituents, such as flavonoids, alkaloids, terpenoids, saponins, tannins, phenolics, and glycosides that have exhibited safe and efficient insulinotropic and antihyperglycemic activities [10].

*A. racemosus*, commonly called Satavar, Shatavari, or Shatamull, is an essential therapeutic plant belonging to the Asparagaceae family that is cultivated in tropical and subtropical India. Its pharmacological applications have been documented in the British and Indian Pharmacopoeias along with conventional medicinal practices [11–13]. This herbal plant is abundant in bioactive constituents including, steroids, flavonoids, saponins, phenolics, and carbohydrate compounds [14]. Previous studies suggest that *A. recemosus* has beneficial effects in neurological disorders, nausea, dysentery, carcinomas, irritation, fatigue, nerve damage, hepatic disease, asthma, breathing difficulties, acidity, and specific bacterial infections [15]. It is also widely utilized as a health stimulant and for managing several clinical complications, including anticancer activities [16], antiulcer [15], antidiarrhoeal [17], immunomodulatory activities [18], antibacterial [19], cardiovascular effects [20], radioprotective agent [21], anti-oxidant [22] and antihyperglycemic [23]. In the previous study *A. racemosus* showed antihyperglycemic impact on streptozotocin induced diabetic rats [23] and also reported to stimulate insulin secretion [24]. Additionally, research suggests that *A. racemosus* exerts inhibitory effects on α-amylase and α-glucosidase enzymes, alongside a high content of bioactive phytochemicals [25] and showed antioxidant properties [26]. Despite these findings, the underlying mechanisms of action remain unclear and require more detailed investigation. The present investigation was designed to further evaluate the antidiabetic potential of the ethanol extract of *A. racemosus* utilizing *in vitro* assays, including starch digestion, glucose diffusion, glucose uptake and DPPH along with phytochemical screening to explore the underlying mechanisms involved in diabetes management.

## 2. Materials and methods

### 2.1. Plant material collection and extract preparation

Roots of *A. racemosus* were procured from Jahangirnagar University in Savar, Dhaka, Bangladesh, botanically verified, and authenticated voucher samples were submitted to the National Herbarium (Bangladesh). The roots were desiccated at 40°C and pulverized into the finest particle (200 mesh) using a cylindrical milling apparatus. Two kilograms powder was obtained via the use of 10 liters of 80% ethanol in a metallic distillation chamber for roughly 4 days at ambient temperature, with the ethanol being replaced every day. The amalgamated extract was subjected to filtration and subsequently vaporized to dryness utilizing an evaporator with a rotary blade. A membrane-based vacuum was employed to expel the extracted material to eliminate the rest of the portion of the solution. The amount of extract (275g) was subsequently frozen-dried utilizing a Varian 801 LY-3-TT freeze drier (Varian, Lexington, MA, USA). The desiccated specimen was preserved at 4°C [23].

### 2.2. *In vitro* starch digestion assay

The starch digestion assay was conducted utilizing a *in vitro* method. Briefly, 100 mg of soluble starch was added to 3 millilitres of purified water in the presence or absence of *A. racemosus* extract across a concentration range of 0.32 to 1000 µg/mL. Acarbose at a similar dosage served as the positive control. The reaction was commenced by introducing 40 µL of thermostable α-amylase from Bacillus licheniformis, with subsequent incubation at 80°C for 20 minutes. The reaction volume was then adjusted to a final volume of 10 millilitres. One milliliter of the mixture was Subsequently added to 0.1 M sodium acetate buffer (pH 4.75) and 30 µL of 0.1% amyloglucosidase obtained from Rhizopus species, and the finished mixture was incubated at 60°C for 30 min to allow complete starch hydrolysis. Following this, the incorporation of the glucose oxidase–peroxidase (GOD–PAP) reagent facilitated the quantification of glucose releases from the specimen utilizing an ELISA assay scanner [27].

### 2.3. *In vitro* glucose diffusion test

An *in vitro* glucose diffusion experiment was conducted utilizing a fiber ester (CE) dialysis filter (MWCO 2000; Spectra/Por®, Spectrum Laboratories, Netherlands) to evaluate glucose movement. The dialysis chamber was filled with 2 mL of 0.9% sodium chloride solution containing 220 mM glucose, either alone or supplemented with varying concentrations of *A. racemosus* extract (40–5000 mg/mL). The extract was evaluated for its ability to enhance solution viscosity and consequently limit glucose movement in the intestinal tract. Both ends of the dialysis bags were securely sealed and immersed in 45 mL of isotonic saline within 50 mL centrifuge tubes. The tubes were maintained at 37°C under continuous gentle agitation with a rotary shaker. At predetermined time intervals, aliquots were withdrawn from the external medium, and glucose levels were measured according to the earlier outlined [28].

### 2.4. *In vitro* glucose uptake assay

Glucose uptake by yeast cells was conducted based on previously prescribed procedure [29]. A 1% (w/v) baking yeast suspension was prepared using purified water and remained at ambient temperature (approximately 25°C) all night. After that, the yeast culture suspension was subsequently centrifuged at 4200 rpm (5 min), and the pellet was washed repeatedly with purified water until a clear supernatant was formed. The final cell suspension was made by diluting the washed yeast to 10% (v/v) with distilled water. Around 1.0 to 5.0 mg/ml of *A. racemosus* plant extracts were combined with DMSO (dimethyl sulfoxide) until they were completely dissolved. Each plant extract mixture was dissolved in 1 mL of glucose solution at various molarities (5, 10, and 25 mM) and preincubated at 37°C for 10 minutes. The interaction process was initiated by introducing 100 µL of the yeast suspension with the combination of glucose and *A. racemosus* ranging from (1.0–5.0 mg/ml), followed by vortexing and then putting it in an incubator for a further 1 hour at 37°C. At the end of the incubation period, the mixtures were rotated and centrifuged at 3800 rpm for 5 min. A spectrophotometer (UV 5100B) was then used to measure the glucose concentration at 520 nm. A control containing all the components except the extract was processed similarly. The percent increase in glucose uptake was determined by the following formula:

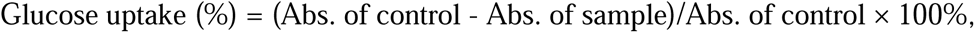

Whereas the control refers to the preparation containing all the ingredients excluding the test sample. Metronidazole was utilized as the reference medicine [30].

### 2.5. *In vitro* DPPH assay

The free radical scavenging ability of the *A. racemosus* extract was evaluated utilizing the 2,2-diphenyl-1-picrylhydrazyl (DPPH) technique. Various concentrations of the *A. racemosus* plant extract (1.6–5000 µg/mL) were prepared in methanol to a final volume of 1 mL. L-ascorbic acid at a similar dosage acted as the reference standard. For each sample, formulated 2 mL of DPPH solution (0.2 mmol/L) was immediately introduced, and the formulations were thoroughly vortexed to ensure uniform mixing. The control solution containing 2 mL of 0.2 mmol/L DPPH solution with 1 mL of distilled water. All the reaction solution were left in the darkness at ambient temperature for 30 min to enable completion of the reaction. The attenuation in absorbance was subsequently measured at 517 nm utilizing an ultraviolet visible spectrophotometric instrument, and the radical scavenging potential was determined relative to the control [31].

### 2.6. Phytochemical screening

*A. racemosus* was screened for preliminary phytochemicals, particularly glycosides, saponins, flavonoids, alkaloids, steroids, tannins, and reducing sugars to find out whether these compounds were present or absent, according to earlier techniques [32–34].

#### Alkaloids

Alkaloid screening was conducted through acidification of 2 millilitres of *A. racemosus* with moderate hydrochloric acid, subsequently introduced 1 millilitre of Dragendroff’s agent. The color transition of the residue from orange to blood red indicated the existence of alkaloids [33].

#### Flavonoids

To test for flavonoids, 4 milliliters of *A. racemosus* were combined with 1.5 milliliters of hot methanol. When two to three drops of hydrochloric acid and magnesium metal were added, the appearance the of mixture turned pink, signifying a positive outcome [33].

#### Tannins

To identify tannins, 10% (w/v) lead acetate solution was added dropwise into 2 millilitres of the *A. racemosus* extract. The appearance of whitish precipitates indicated the existence of tannins [33].

#### Terpenoids

By immersing 1 g of *A. racemosus* in two millilitres of chloroform, 3 ml of strong HCL was subsequently incorporated in a controlled manner to create a distinct phase. The existence of terpenoids was demonstrated by a brownish-red shade at the surface [34].

#### Glycosides

One millilitres of *A. racemosus* was treated with glacial acetic acid containing traces of ferric chloride, followed by the addition of H_2_SO_4_. The formation of a bluish-green hue demonstrated the existence of glycoside [33].

#### Steroids

For steroid identification, 1 ml of *A. racemosus* extract was mixed with 10 ml of chloroform and carefully reacted with 10 ml of H_2_SO_4_. The formation of distinct colored layers, with red in the upper phase and green in the lower phase, was taken as evidence of steroidal constituents [32].

#### Reducing sugars

To identify reducing sugars, 1 ml of *A. racemosus* extract was combined with 1 ml of purified water and a few droplets of Fehling’s reagent, and the solution was heated. The appearance of a brick-red shade confirmed the existence of reducing sugars [33].

### 2.7. Statistical Analysis

All statistical analysis and data interpretation were done using GraphPad Prism 5 (San Diego, CA, USA). Data were analyzed using an unpaired Student’s t-test (two-tailed) and one-way ANOVA followed by Bonferroni post hoc tests. The results were represented as Mean ± SEM with a statistical significance level of *P<*0.05.

## 3. Results

### 3.1. Impact of A. racemosus on starch digestion in vitro

The effects of *A. racemosus* (0.32-1000 ug/mL) and acarbose (0.32-1000 ug/mL) on starch digestion, are illustrated in Fig. 1. *A. racemosus* at concentrations ranging from 200 to 1000 µg/mL significantly inhibited the enzyme-induced glucose digestion of starch, with overall inhibition levels ranging from 14.24% to 21.11% (p<0.05–0.01; Fig 1a) demonstrating a dose-responsive effect in comparison with the untreated control group. Acarbose, used as a reference standard at doses of 1.6 to 1000 µg/mL, demonstrated a concentration-dependent suppression of sugar release from starch, with a range of 34.55% to 70.74% (p<0.01–0.001; Fig. 1b).

**Figure 1:**
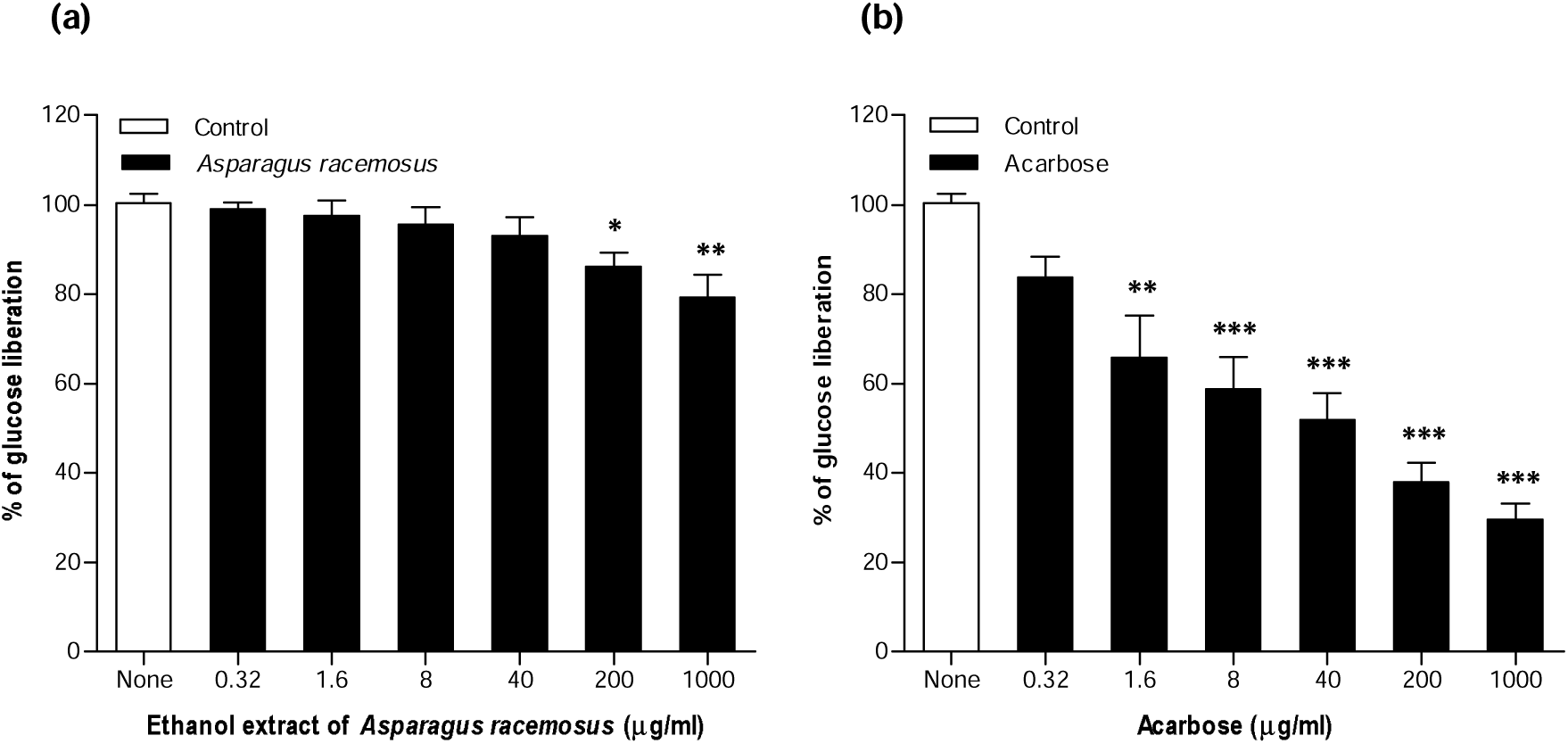
Concentration-dependent effects of *A. racemosus* on *in vitro* carbohydrate digestion; (a) *A. racemosus*, (b) acarbose. The research involved the inclusion and exclusion of *A. racemosus* (0.32–1000 µg/ml) and the positive control acarbose (0.32–1000 µg/ml), subsequent treatment with α-amylase (0.01%) and amyloglucosidase (0.1%). The values are shown as mean ± SEM with n = 4. **\*, **, \*\*\****P<*0.05–*P<*0.001 with respect to the control.

### 3.2. Impact of A. racemosus on glucose diffusion in vitro

The in vitro effects of A. racemosus on glucose movement were evaluated during incubating periods of 0, 3, 6, 12, and 24 hours. A. racemosus significantly reduced the diffusion and absorption of glucose at dosages ranging from 40 to 5,000 µg/ml, in a dose and time-responsive trend. At 0 and 3 hours with A. racemosus, no discernible decrease in glucose permeability was found (Fig. 2a, b). Over 6 and 12 hours of incubation, a significant concentration-responsive decrease (p<0.05–0.001) was noticed, ranging from 9.83% to 15.02% and 9.70% to 16.61% at dosages of 1000 to 5000 µg/mL and 200 to 5000 µg/mL, respectively (Fig. 2c, d). Over 24 hours of treatment, the most significant decrease in glucose movement was observed, ranging from 9.91% to 22.75% at doses of 200–5000 µg/mL (p<0.05–0.001; Fig. 2e).

**Figure 2:**
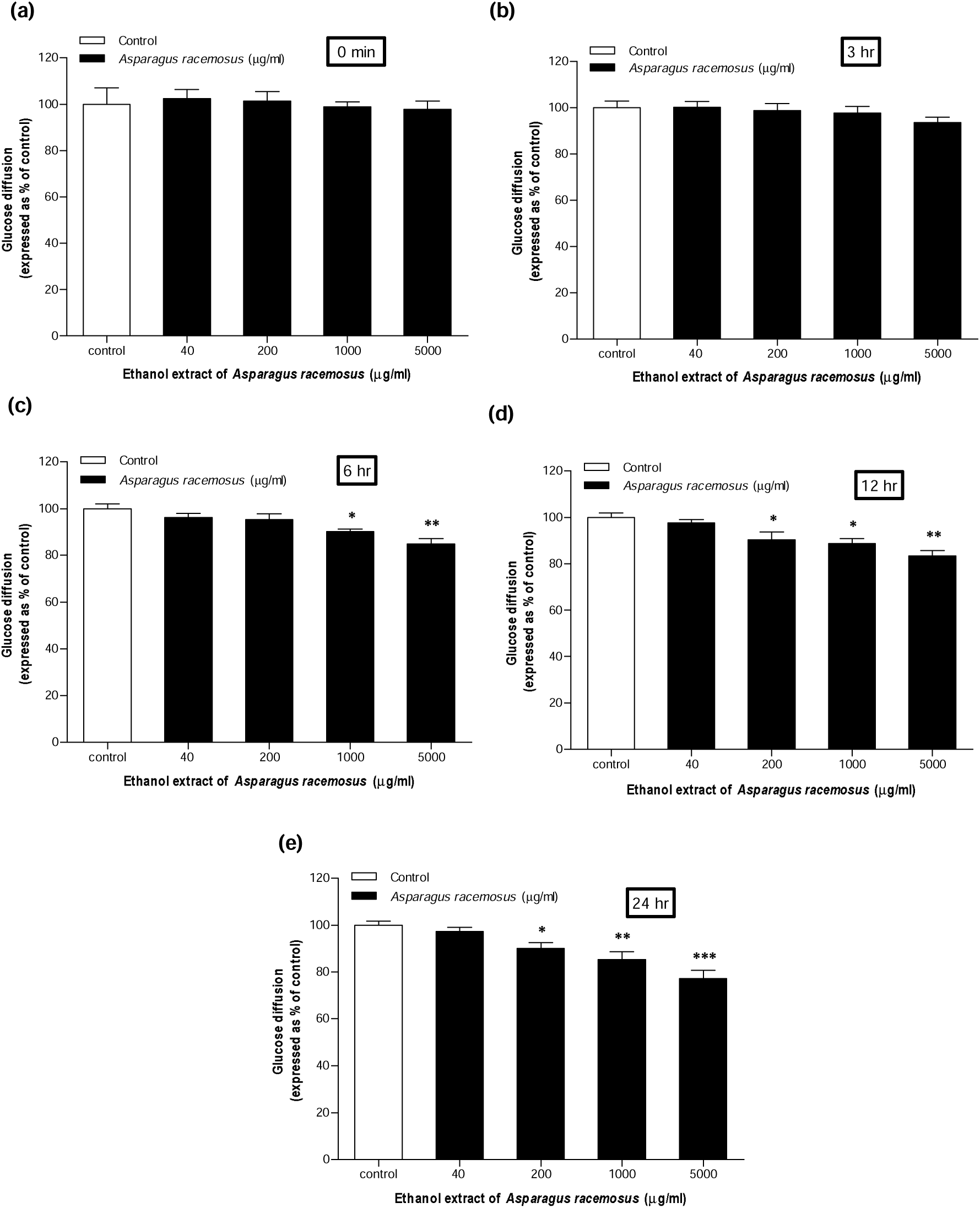
Concentration-dependent effects of *A. racemosus* (a, b, c, d, and e) on *in vitro* glucose diffusion. The investigation was performed with or without *A. racemosus* (40–5000 µg/ml) using cellulose dialysis tubing, and the diffusion of glucose was quantified at (a) 0, (b) 3, (c) 6, (d) 12, and (e) 24 hrs as illustrated by the bars, following the GOD-PAP methodology. The values are shown as the mean ± SEM with n = 4. *, **, *** *P<*0.05–*P<*0.001 with respect to the control.

### 3.3. Impact of A. racemosus on glucose uptake

The ethanolic extract of A. racemosus (2.0–5.0 mg/ml) has shown a significant (p<0.05–0.001) increase in glucose uptake from 45.28% to 65.03% at 5 mM concentrate of glucose in a concentration-responsive manner (Fig. 3a). Also, a notable increase in glucose uptake (p<0.01– 0.001) and (p<0.01–0.001) were detected at 10 mM and 25 mM concentrate of glucose in a concentration-responsive trend, ranging from 34.17% to 47.85% and 11.31% to 16.20%, respectively at A. racemosus (3.0–5.0 and 4.0–5.0 mg/ml) (Fig. 3b, c). Glucose uptake at a beginning dose of 5 mM and 10 mM by the A. racemosus demonstrated comparable activity to the standard drug metronidazole (Fig. 3a, b). However, a significantly higher glucose uptake was observed with metronidazole than A. racemosus at 25 mM glucose (Fig. 3c). Conversely, glucose uptake decreased as the glucose concentration increased compared with that at 5, 10 and 25 mM glucose when the same quantity of A. racemosus was used (Fig. 3a, b, c).

**Figure 3:**
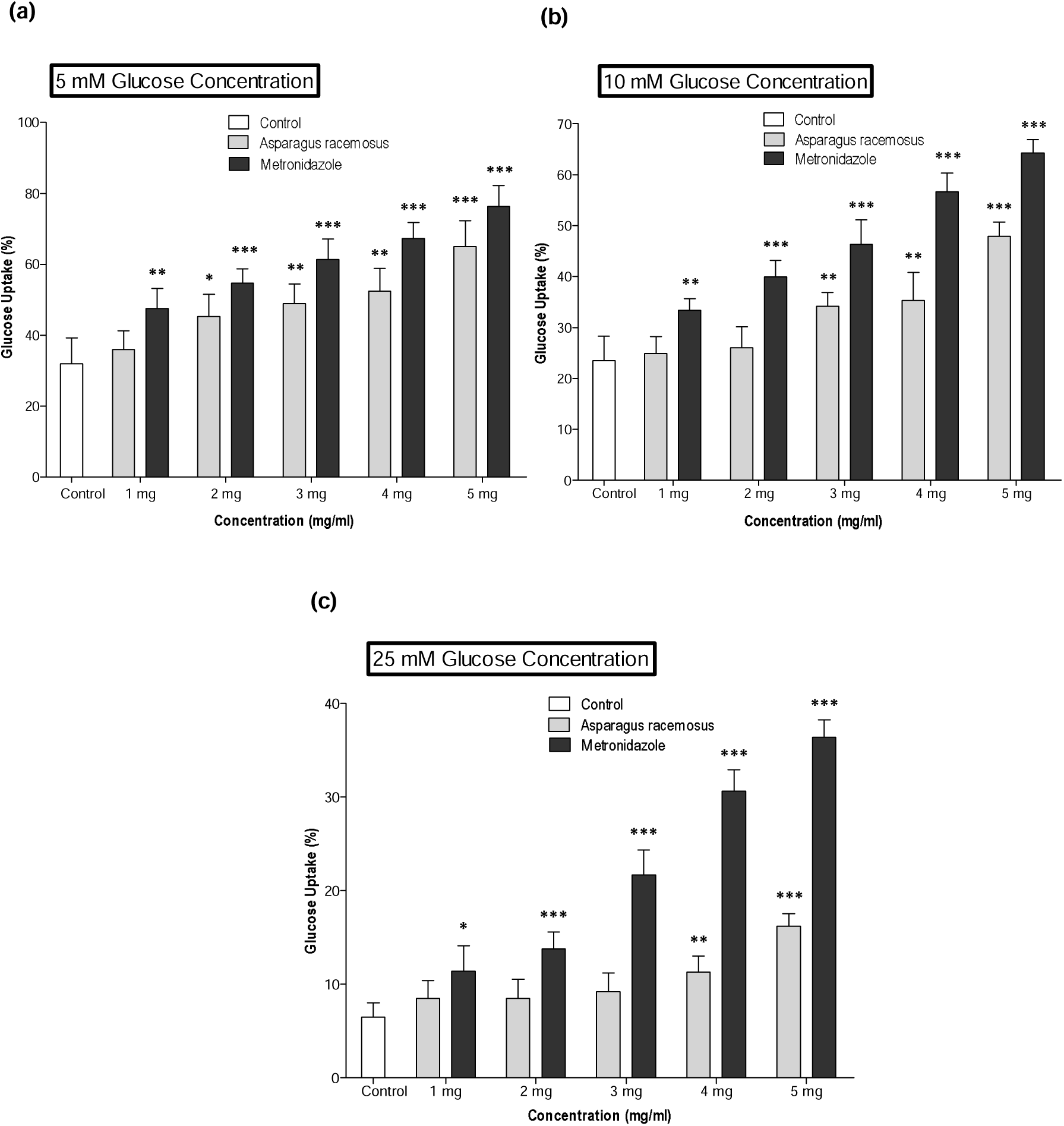
Dose-responsive impacts of *A. racemosus* on *in vitro* glucose uptake capacity; at (a) 5 mM, (b) 10 mM and (c) 25 mM concentration of glucose. The investigation was conducted with or without *A. racemosus* (1.0–5.0 mg/ml) and the positive control metronidazole (1.0–5.0 mg/ml). The values are shown as mean ± SEM with n = 4. *, **, *** *P<*0.05–*P<*0.001 with respect to the control.

### 3.4. Impact of A. racemosus on free radical scavenging

*A. racemosus* significantly inhibited DPPH scavenging capacity by 13.06 ± 3.44 to 42.46 ± 4.62% in a dose-dependent manner at concentrations between 8–5000 µg/mL (p<0.05, 0.001) (Table 1). However, the reference standard, L-ascorbic acid, showed a 13.83 ± 4.08 to 96.47 ± 3.13% (p<0.01–0.001) inhibitory effect on DPPH and increased DPPH free radical scavenging capacity with elevating dosages (1.6–5000 µg/mL), as displayed in Table 1.

**Table 1.**
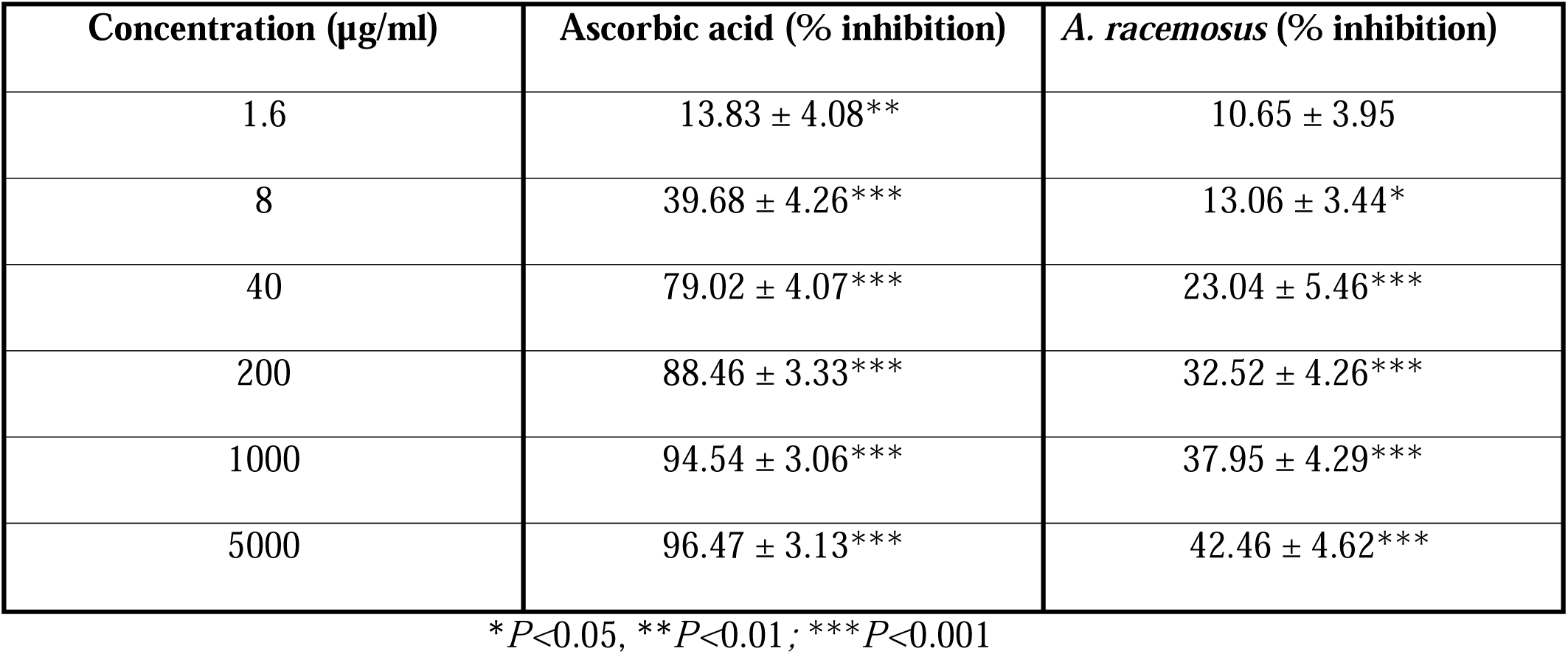
Concentration-responsive impacts of L-ascorbic acid and *A. racemosus* on DPPH scavenging capacity.

| Concentration ($\mu\text{g/mL}$) | Ascorbic acid (% inhibition) | *A. racemosus* (% inhibition) |
| --- | --- | --- |
| 1.6 | $13.83 \pm 4.08^{**}$ | $10.65 \pm 3.95$ |
| 8 | $39.68 \pm 4.26^{***}$ | $13.06 \pm 3.44^{*}$ |
| 40 | $79.02 \pm 4.07^{***}$ | $23.04 \pm 5.46^{***}$ |
| 200 | $88.46 \pm 3.33^{***}$ | $32.52 \pm 4.26^{***}$ |
| 1000 | $94.54 \pm 3.06^{***}$ | $37.95 \pm 4.29^{***}$ |
| 5000 | $96.47 \pm 3.13^{***}$ | $42.46 \pm 4.62^{***}$ |

\* $P < 0.05$ , \*\* $P < 0.01$ ; \*\*\* $P < 0.001$

The experiment was performed using multiple dosages of *A. racemosus* (1.6–5000 µg/ml). Following this, a 30-min incubation with DPPH radical reagent was carried out in atmospheric environments devoid of lighting; the frequency of inhibition of DPPH radicals (%) was measured. The values are shown as mean ± SEM with n = 4; *, **, *** *P<*0.05–*P<*0.001 with respect to the control.

### 3.5. Preliminary phytochemical screening using A. racemosus

A primary phytochemical analysis was performed to find out the existence or lack of potent antidiabetic plant constituents in the ethanol extract of *A. racemosus*. The research confirmed the existence of alkaloids, flavonoids, glycosides, tannins, saponins, steroids and the lack of reducing sugars in *A. racemosus*. (Table 2).

**Table 2.** Preliminary bioactive phytoconstituent screening of the ethanol extract of *A. racemosus*.

| Group | Results |
| --- | --- |
| Alkaloids | + |
| Flavonoids | + |
| Glycoside | + |
| Tannins | + |
| Saponins | + |
| Steroids | + |
| Reducing sugar | - |

(+) = present, (-) = absent of phytoconstituents in *A. racemosus*. Each test was conducted three times (n=3).

## Discussion

*A. racemosus* has reported to show anti-diabetic activities in several studies [23,24,35,36]. However, comprehensive validation and a clear understanding of the underlying mode of action remain limited. The current study aimed to investigate the mechanistic pathway of *A. racemosus* through *in vitro* carbohydrate digestion, glucose diffusion, glucose uptake, DPPH and preliminary phytochemical screening.

An efficient technique for regulating postmeal glycemic levels involves the inhibition of the hydrolysis enzymes α-amylase and α-glucosidase [37]. In the present research, *in vitro* carbohydrate digestion was carried out to verify the effects of *A. racemosus* on α-amylase and α-glucosidase in the breakdown of glucose during the digestion of carbohydrates. The findings of this experiment are consistent with the established enzyme-inhibiting effects of acarbose, the reference standard, as well exhibited a dose-responsive inhibition of starch [38]. Recent studies demonstrated that *A. racemosus* shows inhibitory effects on a-amylase and a-glucosidase and thereby maintains optimal glycemic control [35,36].

Reducing gastrointestinal (GI) absorption and glucose diffusion is a crucial approach for managing blood sugar following dietary intake [39]. Moreover, the diffusion of glucose experiment is an efficient *in vitro* method for estimating the influence of dietary fiber on the retardation of molecular glucose movement in the GI system [40]. The current study investigated the impact of *A. racemosus* on the movement of glucose using a fundamental *in vitro* dialysis method. These results align with an earlier study suggesting that *A. racemosus* might impact glucose absorption [23].

Glucose uptake in yeast cells occurs by facilitating diffusion through the cellular membrane. The glucose content in a designated cell is influenced by multiple variables, including glucose metabolism, glucose transporters, and dietary habits. In response to elevated insulin levels in the circulatory system, lepocytes or myocytes significantly regulate glucose transporting molecules, leading to a reduced blood sugar consequence [30,41]. The results of this research indicate that the ethanolic extract of *A. racemosus* markedly increases glucose uptake in yeast cells, implying enhanced cellular glucose consumption and demonstrating its possible function in blood sugar management. This process may resemble the modes of action of biguanides such as metformin, that lowers glycemic levels by enhancing glucose consumption in tissues around the body [42].

A current investigation revealed that diabetes is linked with oxidative damage and the generation of reactive oxygen species (ROS), leading to impaired insulin action [43]. The present *in vitro* DPPH investigation demonstrated the substantial antioxidant capability of *A. racemosus*, highlighting its ability to scavenge free radicals and mitigate oxidative stress, a crucial aspect of diabetes and its consequences. Prior studies have similarly documented significant DPPH radical scavenging action in *A. racemosus* extracts, and the results of the present study align with earlier findings [44,45].

The phytochemical investigation of the ethanolic extract of *A. racemosus* identified many biologically active substances, including glycosides, saponins, flavonoids, alkaloids, steroids, tannins, and reducing sugars. Alkaloids present in *A. racemosus* are recognized as effective α-glucosidase inhibitors [46, 47]. In addition, tannins have been documented to augment glucose uptake, mitigate inflammation and oxidative stress, and enhance insulin sensitivity [48,49]. Moreover, saponins, especially steroidal saponins, are recognized for their insulin-like properties and their capacity to increase glucose breakdown and weight management via the regulation of adipokine activity [50, 51]. Furthermore, by modulating glucose absorption, insulin signaling, insulin release, and adipose tissue accumulation, flavonoids have been shown to significantly contribute to antihyperglycemic activity [52].

## 5. Conclusion

The current research shows that *A. racemosus* inhibits carbohydrate digestion and absorption, improves glucose uptake, and exerts inhibitory effects on DPPH. These effects may be attributed to the existence of numerous bioactive substances, including glycosides, saponins, flavonoids, alkaloids, steroids and tannins. Consequently, these results demonstrate the efficacy of *A. racemosus* as a beneficial dietary supplement for managing diabetes mellitus. Further research is recommended to isolate and purify the active molecules responsible for the observed antidiabetic effects.

## Acknowledgements

We would like to thank Independent University, Bangladesh (IUB), Dhaka 1229, Bangladesh, for the laboratory support that enabled this research.

## Author contributions

Md. Sifat Rahman and J.M.A. Hannan responsible for the conception and design the study, as well as the supervision of the research. The experiments were conducted and the data were analyzed by Md. Sifat Rahman, Rafia Tasnim, Md. Mashrur Meraj Bhuiyan, Chinmoy Basu, Sakibul Haque Sammo, Bijoy Chandra Sarkar, SM Tajul Islam and Sanjid khan. Md. Sifat Rahman and J.M.A. Hannan also interpreted the findings, created the figures, and wrote the manuscript.

## Declarations

